# Penetration Assessment of Dietary Supplements and Drugs through the Blood-Brain Barrier for Potential Treatment of Parkinson ‘s Disease

**DOI:** 10.1101/362541

**Authors:** Roger Chevalier

## Abstract

Parkinson’s disease (PD) is a progressive neurodegenerative disorder, affecting 2% of the population over the age of 60. To date, there is no disease modifying drugs to prevent dopaminergic neuron loss and abnormal protein deposition in the brains. There is a strong demand for neuroprotective therapies to prevent or slow down dopaminergic neuron degeneration. An essential prerequisite for a compound designed to affect the central nervous system (CNS) is a satisfactory transport through the blood brain barrier (BBB). Numerous posts on the net suggest that both repositioned drugs molecules and active molecules present in dietary supplements may slow down PD’s progression. The logBB is an index of BBB permeability. Starting from quantitative and qualitative permeability data, this study tries to predict logBB values from various physicochemical properties of molecules, including, among others, molecular size, polar surface area (PSA) and logP values. Quantitative logBB models were implemented using MLP (multiple linear regression), PLS (Partial Least Square), AAKR (Auto Associative Kernel Regression) and ECM (Evolving Cluster Method). Qualitative models were carried out with SVM (Support Vector Method).

The paper estimates the BBB permeability of 39 molecules potentially able to slow down PD’s progression and compares the performances of qualitative and quantitative machine learning method used. For information, the current study also gives a short overview of the state of the art on the potential impact of dietary supplements on PD.

## Introduction

Parkinson’s disease (PD) is a progressive neurodegenerative disorder, affecting 2% of the population over the age of 60 [4,5]. PD patients display a loss of dopaminergic neurons in the substantia nigra and the presence of Lewybodies in their brains [4, 5].The current pharmacotherapy for PD patients is limited to symptomatic treatment^1^, which only temporarily reduces motor symptoms but does not prevent neurodegeneration. To date there is no disease modifying drugs to prevent dopaminergic neuron loss and abnormal protein deposition in the brains. There is a strong demand for neuroprotective therapies to prevent or attenuate dopaminergic neuron degeneration.

An essential prerequisite for a pharmaceutical compound designed to affect the central nervous system (CNS) is satisfactory transport through the blood–brain barrier (BBB). The combination of physical (complex tight junctions) and biochemical barriers (metabolic enzyme systems and efflux transporters) make BBB a formidable barrier to candidate CNS drugs and it has been estimated to prevent the brain uptake of more than 98% of potential neurotherapeutics.

In the peculiar case of Parkinson’s disease, a blog managed by Simon Stott [4], aims at bridging the gap between the media headlines and the actual science. It explains the science and research that is currently being conducted on Parkinson’s disease in plain English that even a non scientist will understand. Many posts suggest that numerous molecules absorbed in every day life may slow down Parkinson’s disease. Examples of potential active molecules are present in green tea, coffee, cauliflower, wine etc and can be purchased as dietary supplements (resveratrol, theanine, sulforaphane etc.) An other track refers to applying an existing compound (either an approved drug or a clinical candidate in testing) for one indication to another indication and offers several advantages over traditional drug discovery: reduced cost, risk and time to market.

## Abbreviations

- AUC, Area Under the Curve;
- ADMET, absorption, distribution, metabolism, excretion, toxicity;
- BBB, blood-brain barrier;
- CNS, central nervous system;
- CSF, cerebrospinal fluid;
- DB, Drugbank database;
- MM; Moleculor Mass;
- PC, PubChem database;
- PD; Parkinson’s disease;
- P-glycoprotein; (P-gp), the permeability glycoprotein or plasma glycoprotein
- PSA, Polar Surface Area;
- RMSE, Root Mean Square Error.

## BBB permeability

Before marketing authorization of a candidate CNS drug, systematic validations are essential throughout the whole development process. They consist of simulations and computer-aided engineering resources for in silico testing, rapid prototypes with increasing level of detail for in vitro trials and, (only when working principles and safety have been verified), in vivo trials with animal models [19].

The in silico prediction methods have acquired popularity in the last few decades in the BBB research thanks to their speed, flexibility, low cost and less time-consuming efforts in comparison to in vitro and in vivo approaches. The first pertinent rule of thumb was defined by Lipinski [27] who states that, in general, an orally active drug has no more than one violation of the following criteria:

- No more than 5 hydrogen bond donors (the total number of nitrogen–hydrogen and oxygen–hydrogen bonds
- No more than 10 hydrogen bond acceptors (all nitrogen or oxygen atoms)
- A molecular mass less than 500 daltons
- An octanol-water partition coefficient log P not greater than 5

Lipinski’s criteria have been completed with Veber’s rules which therefore suggest additional filters (rotatable bonds <12 and PSA <140) for drug-likeness:

The rule is applied only to permeation by passive diffusion of orally active drugs through cell membranes. It is a metric that can predict the similarity between two drugs. Thanks to their simplicity, the rules became an efficient tool in the design process.

Drugs that are actively transported through cell membranes by transporter proteins are exceptions to this rule.

Aside from the Lipinski’s rule, a more global approach was elaborated to predict the ability of substances to permeate the BBB successfully. It is the case of logBB, building on experimental data for various drugs like molecules. LogBB is the ratio of the steady-state concentration of the drug molecule in the brain to the concentration in the blood. Predicted logBB parameter was mainly derived from the notion of molecular polar surface area descriptor and octanol-water partition coefficient (logP) to assess compound hydrophobicity and Hbonding capacity (desolvation rate). These two last descriptors were vigorously discussed throughout the literature. For instance, they were implemented in the regression models through many mathematical formulas, such as Clark and Rishton equations:

- logBB = 0.152logP - 0.0148PSA + 0.139
- logBB = 0.155logP - 0.01PSA + 0.164

Unfortunately, experiments to measure logBB are time consuming, laborious and expensive in vitro and even more in vivo. So it is therefore not surprising that the number of published experimental values is limited. Experimental methods aim at assessing:

- In vitro BBB permeability range from artificial membranes (PAMPA) and complex cell culture systems
- In vivo brain uptake index and bolus carotid intravenous.

More generally several mathematical and computational models have been successfully developed to simulate and predict the interaction of compounds with the BBB interface. Two permeability indices, log [brain]/[blood] (logBB) and log permeability-surface area (log PS), are usually retrieved from experimental data of either in vitro or in vivo studies. Thus, modeling algorithms have evolved to quantitatively describe the brain penetration, and to be more precise than the qualitative approach, based on a limited compound classification as either BBB + (a compound that crosses the BBB and attain central nervous system) or BBB - (a compound that does not cross the BBB and does not attain central nervous system).

## Data and Methods

### Construction of the dataset for training and validation

To built our case study, we have analyzed a large number of published papers and identified three free main sources of datasets ([1] [2] [3])

1 - A list of compounds [1] for which logBB values had been measured in vivo, for the most part in rats. This list includes the compounds and the associated descriptors and was collected by (Markus Muehlbacher). As some compounds are not identified both in PubChem and Drug Bank data bases, our list is a subset of list used by [1]. As a reminder, the dataset includes compounds for which the partition had been measured from blood to brain, from plasma to brain and from serum to brain. This set excludes actively transported compounds since their mechanism of passing the BBB is different to those passively entering CNS such as known substrates of P-Gp (bunitrolol, cimetidine, digoxin, domperidone, etoposide, fexofenadine, flunitrazepam, levodopa, loperamide, methotrexate, morphine, nevirapine, phenytoin, quinidine, risperidone, triflupromazine, vincristine, yamatetan).
2 - A table [2] listing FDA-approved drugs (1691) and 3 columns: generic name: Generic name from DrugBank cns drug: TRUE/FALSE, based on whether any of 5 insurance plans checked had the drug listed under “Central Nervous System agents” or similar smiles: isomeric SMILES from DrugBank. When this field is empty it’s usually because the drug is a large biomolecule. <u>Example:</u> Acetaminophen TRUE CC(=O)NC1=CC=C(O)C=C1 Acetazolamide FALSE CC(=O)NC1=NN=C(S1)S(N)(=O)=O Acetic acid FALSE CC([O-])=O This input is described in the web post http://www.Minikel.org/2013/10/04/properties-of-cns-drugs-vs-all-fda-approved-drugs/. It has been collected and analyzed in 2013 by EV Minikel who is on a lifelong quest to develop a treatment or cure for human prion diseases.
3 - Two data bases [3] were used in the study. Data are available on the web: https://github.com/bioinformatics-gao/CASE-BBB-prediction Data Contact: Supplementary information. It was the basic data used by the associated paper publish in 2016 [6]. Datasets are given in the Supplementary Tables S1 and S2. They consists on:

- Drug dataset based on brain/blood concentration ratio (table S1). 213 drugs in total are based on the splitting criteria of Li and coworkers (Li et al., 2005), drugs were divided into BBB+(139 drugs) and BBB‐ (74 drugs) groups according to whether the brain to blood concentration ratio was greater than 0.1 or less than 0.1 respectively (i.e. logBB> ‐1 or < ‐1)
- Drug dataset based on activities in CNS (table S2). Doniger and coworkers composed a drug dataset of 179 BBB+ 145 BBB‐ compounds according to their activities in CNS (Doniger et al., 2002). The compounds which exist in the SIDER database (76 BBB+ and 85 BBB‐ (given in the Supplementary Table S2: Drug Dataset Based on CNS Activities) were used in the framework of our paper as independent test dataset for training and cross validation.

### Construction of the dataset for prediction

As the present study aims at identifying dietary supplements and repositioned drugs which penetrates the BB barrier, three lists of molecules has been collected for analysis (see annex 1):

- Supplements, evoked in the posts described in Simon Stott’s blog [1] or on the net, which may slow down evolution of PD’s according to PD animal or/and cell models.
- Repositioned drugs identified in PD’s blog [1], which may also slow down evolution of PD’s according to PD animal or/and cell models
- Representative CNS drugs aimed at treating brain disorders (depression, Parkinson to decrease symptoms, anxiety etc). This list is built from the French drug dictionary VIDAL and tries to cover the application of CNS marketed drugs.

### Statistical methods

Predicting BBB permeability via statistical modeling has evolved over the past few years, from simple linear regressions to machine learning techniques.

Nevertheless principles of all learning techniques are the same.

1 - First of all an empirical model between drug descriptors and drug BBB permeability (either LogBB value for quantitative model or LogBB True or False for a qualitative model) needs to be tuned during the training phase.
2 - A second dataset is then used to validate the training model by comparing expected and calculated LogBB.
3 - Finally the resulting training model is applied on a third dataset object of the study for the estimation of LogBB. An example is presented in table 3.

**Table 1:**
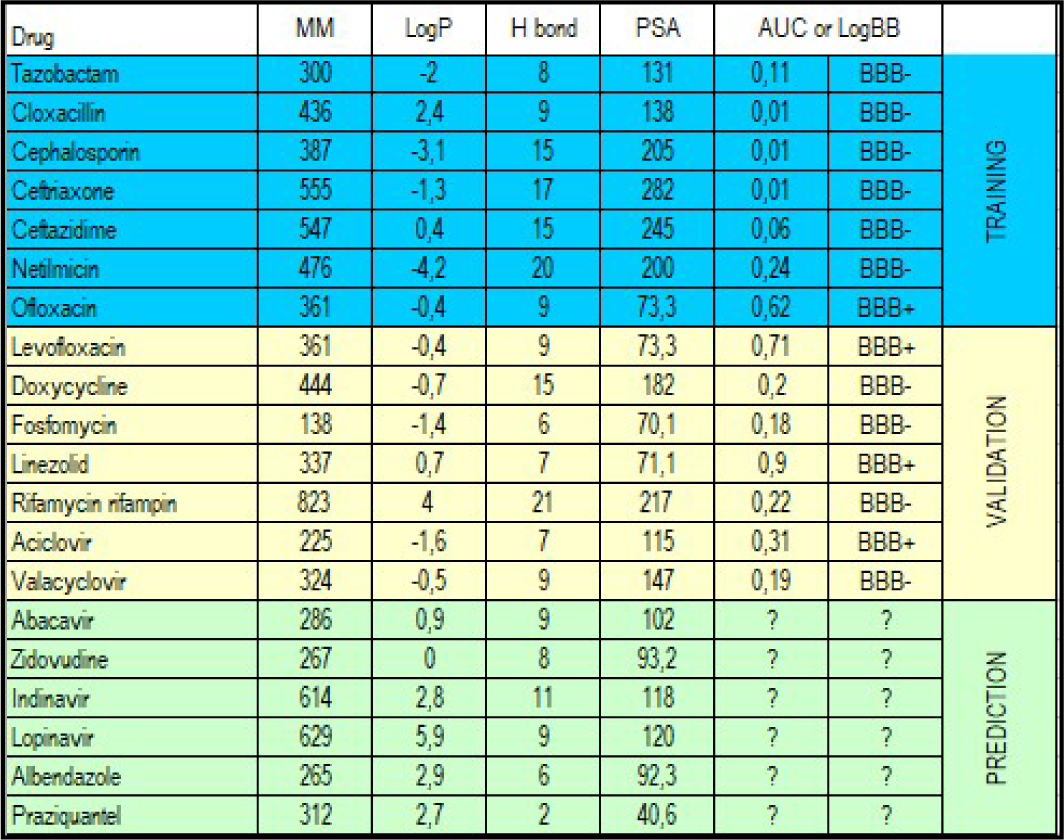
The three steps of a learning technique process.

**Table 2:** Main PubChem representative correlation factors with logBB.

|  | logBB |
| --- | --- |
| COMPLEXITY | -0,2 |
| XLOGP | 0,6 |
| DONOR | -0,6 |
| ACCEPTOR | -0,4 |
| TPSA | -0,7 |

**Table 3:** Main DrugBank representative correlation factors with logBB.

| Properties | logBB |
| --- | --- |
| PSA | -0,6 |
| DB_DONOR_COUNT | -0,5 |
| DB_ACCEPTOR | -0,4 |
| LOGS | -0,2 |
| BIOAVAILABILITY | 0,2 |
| BASICITY | 0,3 |
| LOGP | 0,4 |
| PHYSIOLOGICAL_CHARGE | 0,5 |

**Table 4:** RMSE for training and validation assessed with PC and DB.

|  | PC<br>All 13 | PC<br>6 pp | DB<br>All 19 | DB<br>6 pp |
| --- | --- | --- | --- | --- |
| RMSE (PLS) |  |  |  |  |
| Training mean | 0.57 | 0.6 | 0.46 | 0.59 |
| Validation mean | 0.70 | 0.63 | 0.61 | 0.65 |
| Training std | 0.05 | 0.05 | 0.03 | 0.04 |
| Validation std | 0.15 | 0.05 | 0.04 | 0.05 |

**Table 5:** Number of correct logBB predictions with PC and DB.

|  | PC<br>All 13 | PC<br>6 pp | DB<br>All 19 | DB<br>6 pp |
| --- | --- | --- | --- | --- |
| Number of correct logBB predictions<br>BB+ if error > -0.3 and BB- if error < 0.3 |  |  |  |  |
| Training mean | 64/80 | 62/80 | 67/80 | 62/80 |
| Validation mean | 60/80 | 61/80 | 63/80 | 59/80 |
| Training std | 2.5 | 3.2 | 2.4 | 3 |
| Validation std | 2.5 | 3 | 3 | 3 |

**Table 6:** RMSE for training and validation files calculated with MLR, PLS, AAKR and ECM.

| RMSE | LR | PLS (6) | PLS (12) | AAKR h=1 (2 :8) | AAKR h=0.5 (2 :8) | AAKR h=0.1 (2 :8) | ECM R=0.04 (2 :8) | ECM R=0.02 (2 :8) |
| --- | --- | --- | --- | --- | --- | --- | --- | --- |
| Training mean | 0.457 | 0.458 | 0.438 | 0.23 | 0.06 | 0.01 | 0.11 | 0.04 |
| Validation mean | 0.481 | 0.480 | 0.569 | 0.30 | 0.24 | 0.27 | 0.287 | 0.27 |
| Training std | 0.019 | 0.019 | 0.02 | 0.01 | 0.005 | 0.002 | 0.01 | 0.01 |
| Validation std | 0.020 | 0.020 | 0.15 | 0.02 | 0.02 | 0.02 | 0.025 | 0.027 |

**Table 7:** Mean and standard deviation of the number of exactly predicted compounds [1].

|  | Number of correct prediction Mean |  | Std |
| --- | --- | --- | --- |
| All properties (18) | 150 /185 | 81% | 3.9 |
| MW, logP, Don, Accep, Rotat, PSA | 153/185 | 83% | 4.1 |

**Table 8:** Mean and standard deviation of the number of exactly predicted compounds [2].

|  | Number of correct prediction Mean |  | Std |
| --- | --- | --- | --- |
| All properties (18) | 438 /672 | 65% | 33 |
| MW, logP, Don, Accep, Rotat, PSA | 394/672 | 59% | 46 |

**Table 9:** Percentage of exactly predicted compounds for datasets (Gao S1 and S2)

| Type of data | Mean of the correct prediction number |  | Std of the correct prediction number |
| --- | --- | --- | --- |
| CNS 160 compounds all PC properties (6 to 23) | 58.6 /80 | 73% | 3.7 |
| CNS 160 compounds (PC properties: MW, logP, Donor bonds, Acceptor bonds,, Rotatable bonds, PSA, complexity) | 57.2/80 | 72% | 3.3 |
| AUC (214 compounds) - (PC properties 6 to 23) | 74.6/107 | 70% | 4 |
| AUC (214 compounds) (MW, logP, Donor bonds, Acceptor bonds,, Rotatable bonds, PSA, complexity) | 76.5/107 | 71% | 3.7 |
| CNS Properties selected by Gao [S2] (MW, Volume, surface area, %hydrophilic surf, logP, hydrophilic-lipophilic balance, H-bond donors, H-bond acceptors, 3D H-bonding) | 58/80 | 73% | 3.9 |

**Table 10:** Number of correct logBB predictions for all training datasets.

| Correct logBB predictions in % (validation dataset) | PC Dataset Muehlbaker (1) | DB Minikel (2) | PC Gao (3 CNS) | PC Gao (3 Conc.) |
| --- | --- | --- | --- | --- |
| Validation 6 parameters | 81 % | 65 % | 57 % | 77 % |
| Validation all parameters | 83 % | 59 % | 59 % | 75 % |

**Table 11:** Permeability of selected molecules for PD.

| Supplements | Data [1] AAKR, ECM, MLR, PLS | Score (number of NO) |
| --- | --- | --- |
| Acetylcysteine (NAC) | -0,38 | 0 |
| Cholecalciferol (vitamin D3) | 0,75 | 0 |
| Coenzyme-q10 | 1,16 | 2 |
| Curcumin | 0,05 | 2 |
| Niacin (vitamin B3) | -0,17 | 0 |
| Pyridoxine (vitamin B6) | -0,46 | 0 |
| Resveratrol | -0,05 | 0 |
| Sulforaphane | -0,20 | 0 |
| Thiamine (vitamin B1) | -0,30 | 2 |
| Uric_acid | -0,69 | 2 |

**Table 11: Permeability of selected molecules for PD**
| Repositioned drugs | Data [1] AAKR, ECM, MLR, PLS | Score (number of NO) |
| --- | --- | --- |
| Acetyl_salicylic_acid | -0,02 | 0 |
| Ambroxol | -0,14 | 1 |
| Amlodipine | -0,14 | 3 |
| Bumetanide | -0,36 | 3 |
| Deferiprone | -0,14 | 0 |
| Eliglustat | 0,19 | 2 |
| Ibuprofen | 0,26 | 0 |
| Isradipine | 0,14 | 3 |
| Miglustat | -0,57 | 2 |
| Nifedipine | -0,29 | 3 |
| Nilotinib | 0,29 | 2 |
| Nimodipine | -0,01 | 3 |
| Nitazoxanide | -0,49 | 1 |
| Pioglitazone | 0,08 | 1 |
| Pravastatin | -0,63 | 1 |
| Salbutamol | -0,50 | 3 |
| Simvastatin | 0,29 | 0 |
| Squalamine | -0,68 | 2 |
| Trazodone | 0,29 | 0 |
| Ursodeoxycholicacid | 0,22 | 1 |

Methods used depend on the type of model either qualitative or quantitative. Quantitative models were estimated using MLP (multiple linear regression), PLS (Partial Least Square), AAKR (AutoAssociative Kernel Regression) ([7] [8]) and ECM (Evolving Cluster Method) ([7] [8]). Qualitative models were built by a SVM (Support Vector Method).

MPL, PLS and SDM used are MATLAB algorithms. AAKR and ECM are specific software developed with MATLAB.

## Results and Discussion

### Descriptors

Before starting the study, it is important to identify which physicochemical properties of molecules can explain logBB value. Cross correlation of input data gives a first overview of the explanatory input properties. Two data bases (PubChem and Drug Bank) were exploited to carry out the study. As a consequence, the list of descriptors depends on the properties of the drugs available in the data base.

The only quantitative dataset is the first [1]. 160 drugs out of 371 that are described both in PubChem (13 properties) and Drug Bank (19 properties). The comparison between the two databases shows that two properties (logS and Physiological Charge), are not available in DrugBank. In order to have a better image of the performance of the two databases, 80 compounds were randomly selected for training and the rest for validation with two configurations (all properties or MW, logP, Donor, Acceptor, Rotatable, PSA).

As one can notice, results are very similar for data coming from PC or DB and very good if we think in terms of number of correct predictions.

### Analysis based on Muehlbacher dataset

In brief, our first dataset [1] includes 371 compounds described by 13 descriptors included LogBB. which varies between –2 and 1.64.To generate qualitative models we chose logBB limit as > or <= ‐0.3 between BBB permeable and non permeable

Four data analysis methods were applied to benefit from the fact that logBB is discrete value.

The 371 compounds dataset was split between two data sets of the same dimension for training and validation.

Seven PC descriptors were selected to estimate logBB: complexity, molecular weight, xlogP, donor, acceptor, rotatable bond, TPSA. RMSE was calculated for both training and validation data. H corresponds to the bandwidth coefficient for AAKR and R the maximum radius of the cluster for ECM. The number of Latent Parameters (PLS) is given.

It has to be noted that local estimation methods (validation RMSE mean around 0.27) are twice more performing than linear methods such as MPL and PLS.(RMSE around 0.5).

A quantitative model was created from the 371 compound data set (Y is logBB >-0.3 and N if logBB <=-0.3).

SVM method was applied to estimate logBB status. This model has been created to be able to compare performance of dataset [1] with other datasets ([2] [3]).

Additionally we have compared our coefficients of logBB linear regression with the ones of Clark and Rishton equations [9] [10].

- Clark: logBB = 0.152logP - 0.0148PSA + 0.139
- Rishton: logBB = 0.155logP - 0.01PSA + 0.164

For the complete dataset 371 drugs logBB = 0.127 +0.149logP-0.008PSA and for a more concentrated dataset (274) logBB = 0.1+0.157logP-0.008PSA. Results are consistent with Clark and Rishton findings.

### Analysis applied on Minikel dataset

A table listing FDA-approved drugs (1691) provided by EV Minikel ([2]) which gives a indication of CNS activities based on whether any of 5 insurance plans checked had the drug listed under “Central Nervous System agents”.

As the model is quantitative we used SVM as algorithm. DB was scanned and drugs where numerous properties were excluded. The result consists of 1344 compounds from DB. 672 compounds were randomly selected for training and the rest for validation. The number of correct predictions is of the same order than for study [1] and standard deviation is far better.

Although all the available properties were selected, the ration number of success/ total number of validation compounds is not really satisfying.

### Analysis based on Gao datasets

Two quantitative models were used for this dataset [3]:

- brain/blood concentration ratio (table S1 in [3])
- activities in CNS (table S2 in [3]).

Performance of two datasets (S1 and S2) was assessed with the same approach. Each dataset was split randomly between training and validation file of the same size.

For both CNS and concentration datasets, the rate of correct prediction is rather high and it has to be noted that the performances are of the same level for PC properties and Gao’s properties.

### Robustness of all training models and logBB predictions

First of all, numbers of correct predictions during the validation phase are the largest with datasets [1] and [3 concentration].

All the training models have been applied on dietary supplements, repositioned drugs and CNS marketed drugs for validation of the scope of training datasets. Results are gathered in annex 1. Column 2 presents the average logBB for all modeling methods used (AAKR, ECM, PLS and MLR). As LogBB is the ratio of the steady-state concentration of the drug molecule in the brain to the concentration in the blood, we have defined three prediction zones:

- Ratio <20% LogBB<-0.7 – Red zone - No permeability
- Ratio >=20% and <50% LogBB>=-0.7 and <-0.30 – Yellow zone - Possible permeability
- Ration >= 50% - Green zone - High probability of permeability.

Columns 3, 4 and 5 (annex 1) show results on quantitative models and last column gives the number of No among the processed model‥

Analysis of results suggests the following conclusions:

- Qualitative models [2] [3] are less precise than quantitative model [1];
- Qualitative models seem to be more conservative (several drugs tagged as BBB‐ are in training files);
- List of molecules tagged as permeable is shorter for annex 1 column 3 than for annex 1 column 2.
- All the CNS marketed drugs are identified as permeable both by column 2 and 3 except sulpirid because of low lipid solubility, identified on the net to poorly penetrate the blood-brain barrier.

## Conclusions

The BBB remains a challenge when designing CNS therapeutics, balancing pharmacodynamics with pharmacokinetics, with an approach to be improved. Diverse in silico, in vitro and in vivo methods for the estimation of molecules transferred across the BBB have been devised for pharmaceutical research by industry and academia in the last few decades. Because of the ease of implementation, in silico prediction methods have acquired popularity in the BBB research. Numerous dietary supplements and repositioned drugs have been identified to potentially have an influence on PD evolution. The present study aims at assessing the ability of these molecules to cross the BBB. Using three databases, we defined two type of models for the prediction of logBB value. These models based on logBB prediction are either qualitative or quantitative

To feed our logBB models, we extracted physicochemical properties of molecules from PubChem and DrugBank databases. SVM was privileged to process qualitative model and we used PLS, MLR, AAKR and ECM methods to run quantitative models. From the analysis of results, it has to be noticed that:

- Quantitative models are more precise than qualitative models and furthermore the main problem is that we don’t know the exact meaning of BBB+ and BBB-
- Predicted ADMET features are are less conservative than predictions based on data sets used by the study.
- Models limited to the use of MW, logP, PSA and bonds (donor, acceptance, rotatable) has to be privileged as they appear to be more robust.
- Local estimation (Comparison with nearest neighbors already tagged like ECM, AAKR) is more efficient than interpolation on the overall domain as it can take into account several underlying physicochemical principles such as non linearity.

The list of dietary supplements which are predicted by quantitative and qualitative models and therefore likely to cross the BB are NAC N-acetyl-cysteine, cholecalciferol, niacin, resveratrol and sulforaphane.

Repositioned drugs responding to the same criteria are acid acetyl salicylic acid, deferiprone, ibuprofen, simvastatin and trazodone.

## Acknowledgements

- Simon Stott for the blog scienceofparkinson
- Eric Vallabh Minikel who collected, analyzed and made available a list of FDA-approved drugs with the BBB attribute
- Markus Muehlbacher for collecting, completing and making available a list of compounds for which logBB values had been measured in vivo, for the most part in rats.
- Gao Zhen for making available data used by the associated paper published in 2016. Predict drug permeability to blood–brain-barrier from clinical phenotypes: drug side effects and drug indications
- The author would like to thank Patrick Bernard et Didier et Brigitte Lereverend for their constructive review of the paper.

## Annex 1: Comparison between qualitative and quantitative prediction methods

|  | Quantitative model | Qualitative model |  |  |  |
| --- | --- | --- | --- | --- | --- |
| Drug | Data [1]<br>AAKR,<br>ECM, MLR,<br>PLS | Data [2]<br>SVM | Data [3-conc] SVM | Predicted<br>ADMET features<br>(Drug Bank) | Score of the<br>qualitative models<br>(number of N) |
| Acetylcysteine | -0,38 | Y | Y | not available | 0 |
| Cholecalciferol | 0,75 | Y | Y | BBB+ 0,96 | 0 |
| Coenzyme-q10 | 1,16 | N | N | not available | 2 |
| Curcumin | 0,05 | N | N | not available | 2 |
| Egcg | -1,16 | N | N | not available | 2 |
| Glutathione | -1,57 | N | N | BBB+ 0,65 | 3 |
| Inosine | -1,00 | N | Y | BBB+ 0,80 | 2 |
| Mannitol | -1,48 | N | Y | BBB- 0,60 | 2 |
| Niacin | -0,17 | Y | Y | BBB+ 0,96 | 0 |
| Pyridoxine | -0,46 | Y | Y | BBB+ 0,69 | 0 |
| Resveratrol | -0,05 | Y | Y | BBB+ 0,59 | 0 |
| Sulforaphane | -0,20 | Y | Y | not available | 0 |
| Theanine | -1,00 | N | N | not available | 3 |
| Thiamine | -0,30 | N | N | BBB+ 0,93 | 2 |
| Uric_acid | -0,69 | N | N | not available | 2 |
| Uridine | -0,96 | N | Y | BBB+ 0,77 | 2 |
| Drug | Data [1]<br>AAKR,<br>ECM, MLR,<br>PLS | Data [2]<br>SVM | Data [3-conc] SVM | Predicted<br>ADMET features<br>(Drug Bank) | Score of the<br>qualitative models<br>(number of N) |
| Acetyl_salicylic_acid | -0,02 | Y | Y | BBB+ 0,94 | 0 |
| Ambroxol | -0,14 | Y | N | not available | 1 |
| Amlodipine | -0,14 | N | N | BBB- 0,77 | 3 |
| Bumetanide | -0,36 | N | N | BBB- 0,66 | 3 |
| Deferiprone | -0,14 | Y | Y | BBB+ 0,94 | 0 |
| Doxycycline | -1,26 | N | N | BBB- 0,99 | 4 |
| Eliglustat | 0,19 | N | N | not available | 2 |
| Ibuprofen | 0,26 | Y | Y | BBB+ 0,96 | 0 |
| Isradipine | 0,14 | N | N | BBB- 0,84 | 3 |
| Leucovorin | -1,43 | N | N | BBB- 0,78 | 4 |
| Miglustat | -0,57 | N | Y | BBB- 0,90 | 2 |
| Nifedipine | -0,29 | N | N | BBB- 0,95 | 3 |
| Nilotinib | 0,29 | N | N | BBB+ 0,90 | 2 |
| Nimodipine | -0,01 | N | N | BBB- 0,93 | 3 |
| Nitazoxanide | -0,49 | N | Y | BBB+ 0,72 | 1 |
| Pioglitazone | 0,08 | Y | N | BBB+ 0,88 | 1 |
| Pravastatin | -0,63 | N | Y | not available | 1 |
| Salbutamol | -0,50 | N | N | BBB- 0,96 | 3 |
| Simvastatin | 0,29 | Y | Y | BBB+ 0,94 | 0 |
| Squalamine | -0,68 | N | N | not available | 2 |
| Trazodone | 0,29 | Y | Y | BBB+ 0,98 | 0 |
| Trehalose | -1,64 | N | Y | not available | 2 |
| Ursodeoxycholicacid | 0,22 | N | Y | BBB+ 0,93 | 1 |
| Drug | Data [1]<br>AAKR,<br>ECM, MLR,<br>PLS | Data [2]<br>SVM | Data [3-conc] SVM | Predicted<br>ADMET features<br>(Drug Bank) | Score of the<br>quaitative models<br>(number of N) |
| Agomelatine | 0,16 | Y | Y | BBB+ 0,99 | 0 |
| Alprazolam | 0,15 | Y | Y | BBB+ 0,98 | 0 |
| Amantadine | 0,16 | Y | Y | BBB+ 0,98 | 0 |
| Amoxapine | 0,11 | Y | Y | BBB+ 0,98 | 0 |
| Aripiprazole | 0,38 | Y | N | BBB+ 0,99 | 1 |
| Citalopram | 0,41 | Y | Y | BBB+ 0,97 | 0 |
| Clomipramine | 0,75 | Y | Y | BBB+ 0,98 | 0 |
| Diazepam | 0,28 | Y | Y | BBB+ 0,99 | 0 |
| Doxepin | 0,61 | Y | Y | BBB+ 0,94 | 0 |
| Droperidol | 0,25 | Y | N | BBB+ 0,96 | 1 |
| Duloxetine | 0,35 | Y | Y | BBB+ 0,98 | 0 |
| Fluoxetine | 0,65 | Y | Y | BBB+ 0,98 | 0 |
| Isoniazid | -0,42 | Y | Y | BBB+ 0,99 | 0 |
| Loxapine | 0,32 | Y | Y | BBB+ 0,98 | 0 |
| Nortriptyline | 0,51 | Y | Y | BBB+ 0,95 | 0 |
| Pimozide | 0,54 | Y | N | BBB+ 0,97 | 1 |
| Rasagiline (PD) | 0,21 | Y | Y | BBB+ 0,98 | 0 |
| Rotigotine (PD) | 0,42 | Y | Y | BBB+ 0,98 | 0 |
| Selegiline (PD) | 0,43 | Y | Y | BBB+ 0,98 | 0 |
| Sertraline | 0,65 | Y | Y | BBB+ 0,97 | 0 |
| Sulpiride | -0,52 | N | N | BBB+ 0,94 | 2 |
| Zolpidem | 0,04 | Y | Y | BBB+ 0,94 | 0 |

## Potential impact of dietary supplements on PD

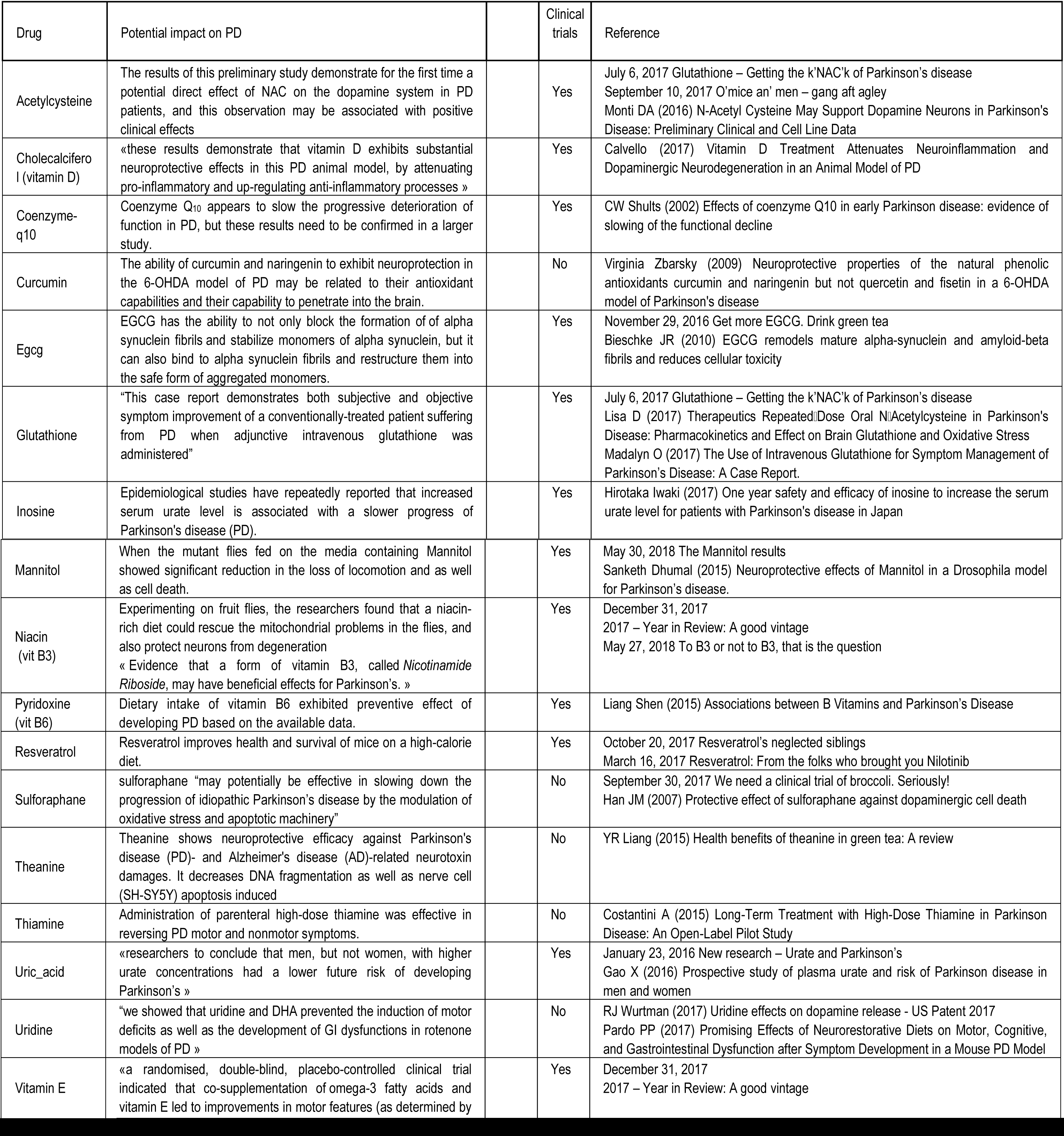

## List of descriptors from PubChem and DrugBank databases

The overall list of descriptors for PubChem is:

|  |  |
| --- | --- |
| CACTVS_COMPLEXITY | TOTAL_CHARGE |
| CACTVS_HBOND_ACCEPTOR | HEAVY_ATOM_COUNT |
| CACTVS_HBOND_DONOR | ATOM_DEF_STEREO_COUNT |
| CACTVS_ROTATABLE_BOND | ATOM_UDEF_STEREO_COUNT |
| XLOGP3 | BOND_DEF_STEREO_COUNT |
| EXACT_MASS | BOND_UDEF_STEREO_COUNT |
| MOLECULAR_WEIGHT | ISOTOPIC_ATOM_COUNT |
| CACTVS_TPSA | COMPONENT_COUNT |
| MONOISOTOPIC_WEIGHT | CACTVS_TAUTO_COUNT |

And for Drug Bank

|  |  |
| --- | --- |
| MOLECULAR_WEIGHT | JCHEM_MDDR_LIKE_RULE |
| EXACT_MASS | JCHEM_NUMBER_OF_RINGS |
| JCHEM_ACCEPTOR_COUNT | JCHEM_PHYSIOLOGICAL_CHARGE |
| JCHEM_ATOM_COUNT | JCHEM_PKA |
| JCHEM_AVERAGE_POLARIZABILITY | JCHEM_PKA_STRONGEST_ACIDIC |
| JCHEM_BIOAVAILABILITY | JCHEM_PKA_STRONGEST_BASIC |
| JCHEM_DONOR_COUNT | JCHEM_POLAR_SURFACE_AREA |
| JCHEM_FORMAL_CHARGE | JCHEM_REFRACTIVITY |
| JCHEM_GHOSE_FILTER | JCHEM_ROTATABLE_BOND_COUNT |
| ALOGPS_LOGP | JCHEM_RULE_OF_FIVE |
| JCHEM_LOGP | ALOGPS_SOLUBILITY |
| ALOGPS_LOGS | JCHEM_VEBER_RULE |

## Footnotes

1 Dopamine in blood is unable to cross the blood-brain barrier to reach the brain. In Parkinson’s disease and doparesponsive dystonia there is a deficiency of dopamine in specific areas of the brain like the basal ganglia. Levodopa is a dopamine precursor. It enters the brain by crossing the blood-brain barrier. It is typically co-administered with an inhibitor of peripheral decarboxylation (DDC, dopa decarboxylase), such as carbidopa or benserazide. This additional drug prevents breakdown of the levodopa to dopamine in the peripheral blood and ensures that maximum amount reaches the brain (www.news-medical.net/health/Dopamine-Therapeutic-Use.aspx). As an example, several clinical trials based on repositioned drugs (ambroxol, isradipin, simvastatin etc) are launched in the case of Parkinson’s. All these tests bring a lot of hope among Parkinson’s community as this condition is considered to be incurable. Consequently as it can be perceived on social networks, patients do not hesitate to take dietary supplements or marketed drugs to slow down PD without knowing if these molecules can cross BBB. The present study relies on several published data and papers aiming to assess the influence of the drugs’ physicochemical properties on the entry of Parkinson’s potential dietary supplements and repositioned drugs into the cerebrospinal fluid (CSF).

